# Characterization of the biofilm landscape of *Bacillus subtilis* by spatial microproteomics

**DOI:** 10.64898/2026.02.02.702398

**Authors:** Kevin J. Zemaitis, Mowei Zhou, Sarah M. Yannarell, James M. Fulcher, Arunima Bhattacharjee, Marija Veličković, David J. Degnan, Elizabeth A. Shank, Christopher R. Anderton, William Kew, Ljiljana Paša-Tolić, Dušan Veličković

**Author notes:** **CORRESPONDING AUTHOR(S):** Kevin J. Zemaitis –, Dusan Veličković –. Department of Chemistry, Zhejiang University, Hangzhou, Zhejiang 310058, China (Mowei Zhou).

## Abstract

Bulk proteomics has been demonstrated to differentiate subpopulations based on molecular phenotypes within bacterial colonies, yet advanced analyses by mass spectrometry imaging (MSI) hold even greater promise for the future. This technology can enable high-throughput spatial phenotyping that can directly visualize distinct components of various biomolecular mechanisms with high mass resolving power in high spatial resolution analyses. Here, we applied MSI for intact protein imaging directly from thin cross sections of a biofilm of *Bacillus subtilis* and after minimal preparation we detected more than 285 unique isotopic envelopes corresponding to unique proteoforms. We paired our MSI analyses with bulk top-down proteomics (TDP) to form extensive experimental libraries, which provided us with high confidence MSI annotations based upon isotopic matching to validated post-translational modifications (PTMs) and truncations. This joint application of MSI and TDP allowed us to describe the microscal e spatial proteomic landscape within the *B. subtilis* biofilm. This study further demonstrated the feasibility of detecting differentiated subpopulations of cells through the identification of proteoforms of cannibalistic protein toxins as well as those involved in active sporulation to highly localized areas within the central and outermost periphery of the biofilm.

**GRAPHICAL ABSTRACT:** 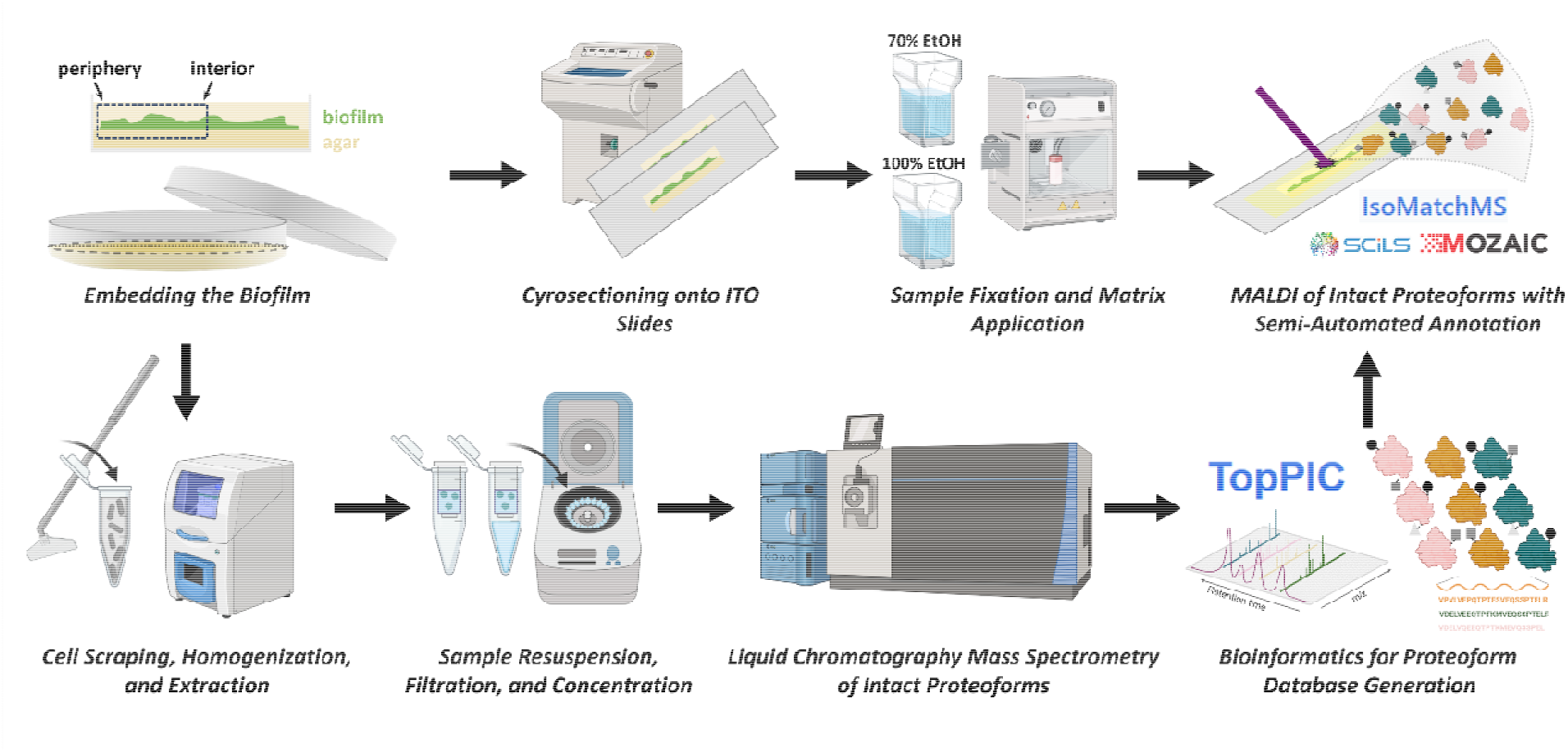

## INTRODUCTION

*Bacillus subtilis* biofilms form distinct spatial microenvironments where unique biological functions such as sporulation, colonization, and metabolite production are governed by underlying mechanisms.(1, 2) The proteins carrying out these biochemical mechanisms can be used as proxies to localize cellular phenotypes from the central core to the outer periphery of a colony.(3) Not only are these subpopulations spatiotemporally dynamic, but they directly result from differential genetic expression,(4) with mutations driving further heterogeneous responses.(5) Using *B. subtilis* as a model organism for biofilm development,(6) we conducted top-down proteomics (TDP) and matrix-assisted laser desorption/ionization (MALDI) mass spectrometry imaging (MSI) on thin sections of the biofilm to explore the intact protein landscape revealing differentiated subpopulations.(7, 8) MSI has been broadly applied within microbiology for the detection of primary and secondary metabolites and highly abundant lipids.(9) The detection of intact proteins distributions hold unique promise,(10) where species, subpopulations, and molecular phenotypes can be identified by differentially abundant proteins associated with unique biochemical mechanisms in a manner similar to spatial biotyping.(11)

Relatedstudies have focused on MALDI-MSI of small peptide toxins within *B. subtilis* (e.g., sporulation delaying protein, SDP; sporulation killing factor, SKF; and epipeptide, EPE),(12, 13) which traced cannibalistic cell-types in wild-type and knockout mutants. In addition to MALDI-MSI mapping the localization of these cannibalistic toxins,(12) further insight has been drawn into the function and interdependence of these peptide toxins in the control of sporulation and further biofilm development through genetic knockouts.(14) Thus, MALDI-MSI is an enabling tool for microbiology, enabling the direct localization of lipids, primary and secondary metabolites, and peptide toxins, which can enhance more traditional spectroscopy completed on cultured biofilms. Here, we demonstrate discovery-based efforts for contextualization of spatial dependencies of intercellular and extracellular proteins(15) providing the location of larger abundant proteomic mechanisms responsible for cellular differentiation and biofilm production.(10)

## METHODOLOGY

### Sample Preparation of B. subtilis for Top-Down Proteomics (TDP)

For TDP, *B. subtilis* was cultured similarly to as previously described on MSgg agar plates.(1) After 48 hours of growth isolated *B. subtilis* colonies were scraped with a cell scraper and collected in Eppendorf tubes. These tubes were subsequently frozen and lyophilized. The lyophilized sample was bead beaten with a 3 mm tungsten carbide beads at 1,400⍰c.p.m. for 2⍰min using a HG-600 Geno/Grinder 2010 Tissue Homogenizer and Cell Lyser (Cole Palmer, Metuchen, NJ), using cooled cryoblocks to inhibit sample heating. The powdered sample was resuspended in Type 1 water and cold (™20 °C) chloroform:methanol mixture (2:1, *v/v*) in chloroform-compatible 2 mL Sorenson MulTI SafeSeal microcentrifuge tubes (Sorenson Bioscience, Salt Lake City, UT). The sample was vortexed for 1⍰min, followed by incubation on ice for 5⍰min, then vortexed again for 1⍰min and centrifuged at 10,000 g at 4⍰°C for 10⍰min. The top and bottom phases were discarded. The remaining protein interlayer was washed with cold 100% methanol, vortexed, and centrifuged for 5 min to pellet the protein. Supernatant was discarded. The protein interlayer was resuspended in 8 M urea prepared in 50 mM ammonium bicarbonate (pH 8). Samples were subsequently cleaned and concentrated using Amicon ultra centrifugal filtration molecular weight cutoff (MWCO) filter tubes (Millipore, Bedford, MA) following the manufacturers standard operating procedure for centrifuging at 14,000 g. Filtration with a 100 kDa filter resulted in concentrated protein that was subsequently transferred to a 10 kDa MWCO filter for desalting. After filtration the protein concentration was determined using a bicinchoninic acid (BCA) assay (Thermo Scientific, Waltham, MA) and samples were diluted to 0.5 µg/µL in water with 0.1% formic acid.

### Liquid Chromatography Mass Spectrometry for Top-Down Proteomics

Liquid chromatography mass spectrometry (LC-MS) was completed on both an Exploris 480 Orbitrap MS (Thermo Scientific, Bremen, DE) and a Lumos Tribrid Orbitrap MS (Thermo Scientific, San Jose, CA) injecting protein onto a NanoAcquity (Waters, Millford, MA) with dual pumps configuration; use of multiple instruments was for validation and is not required for successful TDP. Binary solvents for mobile phase (MP) A and B were 0.2% formic acid in water (MPA), and 0.2% formic acid in acetonitrile (MPB) for both pumps. The stationary phase material for in-house packed reversed phase LC columns was C2 particle (SMTC2MEB2-3-300, Separation Methods Technologies, Newark, DE). 10 μl injections were loaded onto a short trap column (150 µm i.d., 5 cm long) and desalted with a flow of 3 μl/min with 5% MPB for 5 min. Then the trap column was switched to the analytical column (100 µm i.d., 50 cm long) with flow rate of 0.3 μl/min. Separation gradient started at 5% MPB to 15% MPB over 1 minute and was linearly ramped to 90% MPB over 89 min.

The Exploris 480 Orbitrap MS (Thermo Scientific, Bremen, DE) was operated with the following source parameters including a spray voltage of 2.2 kV, transfer capillary temperature of 275 °C°, and ion funnel RF amplitude set to 60%. The instrument was set to “intact protein” application mode with standard pressures, and data was collected as full profile. MS1 acquisitions were collected with 5 microscans at a mass resolution setting of 120k over a scan range of *m/z* 500 to 2000 with a normalized AGC target of 400%. Data dependent MS2 scans were collected with 2 microscans over a scan range of *m/z* 300 to 2000 with a normalized AGC target of 1000%. MS2 used a dynamic exclusion after 1 time over a duration of 40s with an isolation window of ±2 *m/z* with fragmentation by HCD at a fixed normalized collision energy of 35%.

The Lumos Tribrid Orbitrap MS (Thermo Scientific, San Jose, CA) was operated with the following source parameters including a spray voltage of 2.2 kV, transfer capillary temperature of 275 °C, and ion funnel RF amplitude set to 50%. The instrument was set to “peptide” application mode with standard pressures, and data was collected as full profile. MS1 acquisitions were collected with 3 microscans at a mass resolution setting of 120k over a scan range of *m/z* 400 to 2000 with a normalized AGC target of 200%. Data dependent MS2 scans were collected with 1 microscans over a scan range of *m/z* 300 to 2000 with a normalized AGC target of 500% and maximum injection time of 118 ms. MS2 used a dynamic exclusion after 1 time over a duration of 60s with an isolation window of ±1 *m/z* with fragmentation by HCD at a stepped collision energy of 20V, 30V, and 40V.

### Top-down proteomics data analysis

MS instrument “.RAW” files were first converted into “.mzML” format with MSConvert before processing with the open-search software TopPIC Suite (version 1.4.13).(16) For TopFD deconvolution, a precursor window of 1 and 2 *m/z* corresponding to the Lumos and Exploris respectively, error tolerance of 0.02 *m/z*, MS1 signal to noise ratio (SNR) of 3, MS2 SNR of 1, maximum charge of 30, and maximum monoisotopic mass of 50,000 Da were applied. Specific settings for the TopPIC search included an error tolerance of 15 ppm, PrSM cluster tolerance of 3.2 Da, FDR of 0.01, and a maximum and minimum unknown mass shift of ±500 Da, and a maximum of one unknown mass shift.

### Producing Thin Bacterial Biofilms for Imaging

For MALDI-MSI *B. subtilis* colonies were cultured on MSgg growth medium as previously described,(2) and a thin sectioning protocol was applied as described in greater depth elsewhere.(17, 18) Briefly, biofilm agar blocks were quartered and transferred to molds (22-363-553, Fisher Scientific). These molds were then snap frozen at -80 °C, overlaid with 4% (wt/vol) agarose (Lonza Bioscience) and frozen at -80 °C for 15⍰min. Sectioning was completed as previously reported using a Cryostar NX70 cryotome (Thermo Scientific, Runcorn, UK) and cross-sections were taken at 20 µm thickness onto ITO slides (Bruker Daltonics, Bremen, DE). These sections were stored at -80 °C until analysis.

### Preparation for Intact Proteoform Imaging

Slides were prepared for MALDI by vacuum desiccation at -15 inHg for 30 min with subsequent washing in a two-part series for prior to matrix deposition. Washing was completed with submersion in a solution of 70% ethanol for 30 sec, then submersion in a solution of 100% ethanol for 30 sec. After this, the biofilms were dried by a gentle stream of nitrogen for 30 sec. A M5 Sprayer (HTX Technologies, Chapel Hill, NC) was then used to spray the surface with 5 crisscross (CC) passes of a solution of 5% acetic acid (v/v) in 50% ethanol with a flow rate of 150 μL/min, a nozzle temperature of 30.0 °C, and the velocity set to 1,250 mm/min with 10 PSI of nitrogen gas to boost proteoform signal intensity.(19) A 5 s drying period was added between each of the passes with a 3 mm track spacing, the same sprayer was then immediately used to deposit 15 mg/mL 2,5-DHA in 90% acetonitrile with 0.2% TFA. Only the supernatant of the dissolved matrix was sprayed after sonication and centrifugation. The flow rate of the matrix was 150 μL/min with a nozzle temperature of 30.0 °C, and the velocity was set to 1,300 mm/min with 10 PSI of nitrogen gas. The matrix was applied with a 2 mm track spacing and a crisscross pattern, and the matrix coverage was calculated to be 277 μg/cm^2^. The matrix was then recrystallized with 5% acetic acid solution (v/v) in water at 38.5 °C and dried for 3.5 min, using an apparatus similar to that previously reported.(20)

### MALDI Mass Spectrometry Imaging of Sectioned Biofilms

An elevated pressure (EP) MALDI source with an atmospheric pressure ionization inlet (Spectroglyph, LLC, Kennewick, WA) was mounted on a research grade Q Exactive HF Orbitrap MS (Thermo Scientific, Bremen, DE); this instrument was upgraded with ultrahigh mass range (UHMR) boards and operated under custom privileges licenses. This platform is described in greater depth elsewhere,(21). Briefly, the MALDI source was operated at 7.8 Torr, with fore vacuum of the spectrometer at 1.0 mbar, and an ultrahigh vacuum adjusted to 2.54e-10 mbar with argon gas. Spectra were acquired over a *m/z* range of 2,000 to 20,000, and acquisitions were carried out with preset resolution of 240k, resulting in a transient of 512 ms, and observed mass resolving power of 45k at *m/z* 6,725. The laser diode of the Explorer One 349 nm Nd:YLF (Spectra-Physics, Milpitas, California) was set to output an average of 1.1 µJ/pulse at 1 kHz, and ion were accumulated for 500 ms – resulting in 500 laser shots per pixel. Imaging was carried out at 20 μm spatial resolution, and the resultant desorption of the crater measured by light microscopy was found to be roughly 12 μm × 15 μm, which did not oversample the surface.

### Mass Spectrometry Imaging Data Analysis

Visualization was completed with automatic import of the .RAW acquisition file and .xml of the pixel coordinates into SCiLS Lab Pro (v.2021c, Bruker Daltonics, Bremen, DE). Ion images were generated by extracting singular isotopic peaks for the proteoforms after root mean square normalization (RMS). For proteoform assignments, an experimental library of proteoforms from TDP LC-MS/MS were used as custom databases and isotopic profiles were manually matched and mass shifts annotated. Mozaic (v.2023.4.0.b3, Spectroswiss, Lausanne, CH) was used to average the 34,698 acquisitions within the MALDI-MSI experiment, after averaging the spectra was baseline corrected using the model-free method with a median size of 250, and other default parameters for number of points, sigma, iterations and level (11, 5, 1, 1). A developed package IsoMatchMS, was used to generate candidate matches based on isotopic distribution matching; this package is explained in greater depth elsewhere.(22) Briefly, an entire average of the acquisition file was exported as a .mzML file after processing. This file was imported with annotations and accurate masses with <5 ppm error were considered, and further filtering of the matches occurred with Pearson correlation score of ≥0.7. Further manual verification of annotated and non-annotated proteoforms were searched against multiple adducts.

## RESULTS AND DISCUSSION

Here, we use spatial proteomics to provide direct insights into differentiated bacterial cell subpopulations. Previous analyses of these samples had identified distinct distributions of lipids and surfactants,(2) while here an optimized sample preparation that included mild ethanol fixation enabled the detection of intact protein distributions **(Figure 1a)**. We found that further desalting was not necessary, unlike within mammalian and plant tissues previously analyzed on this platform, or for colonies dried directly onto ITO slides and analyzed from the top down (23). Analyses of these samples were enabled by coupling a MALDI source to a research-grade UHMR HF Orbitrap MS;(19) this combination yields rich spectra with high intensities of intact proteins and proteoforms up to 20 kDa in mass **(Figure 1r)**, although the largest detected species from the biofilm was roughly 13.5 kDa.(24) Subsequently, we performed semi-automated annotation and evaluation of the proteomic spatial heterogeneity within the biofilm. Overall, more than 285 unique isotopic envelopes from proteoforms were detected **(Figure 1r)** and could be annotated via UniProt or experimentally generated TDP libraries – enabling precise annotation of PTMs and truncations.(25) Manual validation was performed after isotopic matching,(22) and sample-specific TDP libraries were imperative for the successful annotation of ion images.

**Figure 1.**
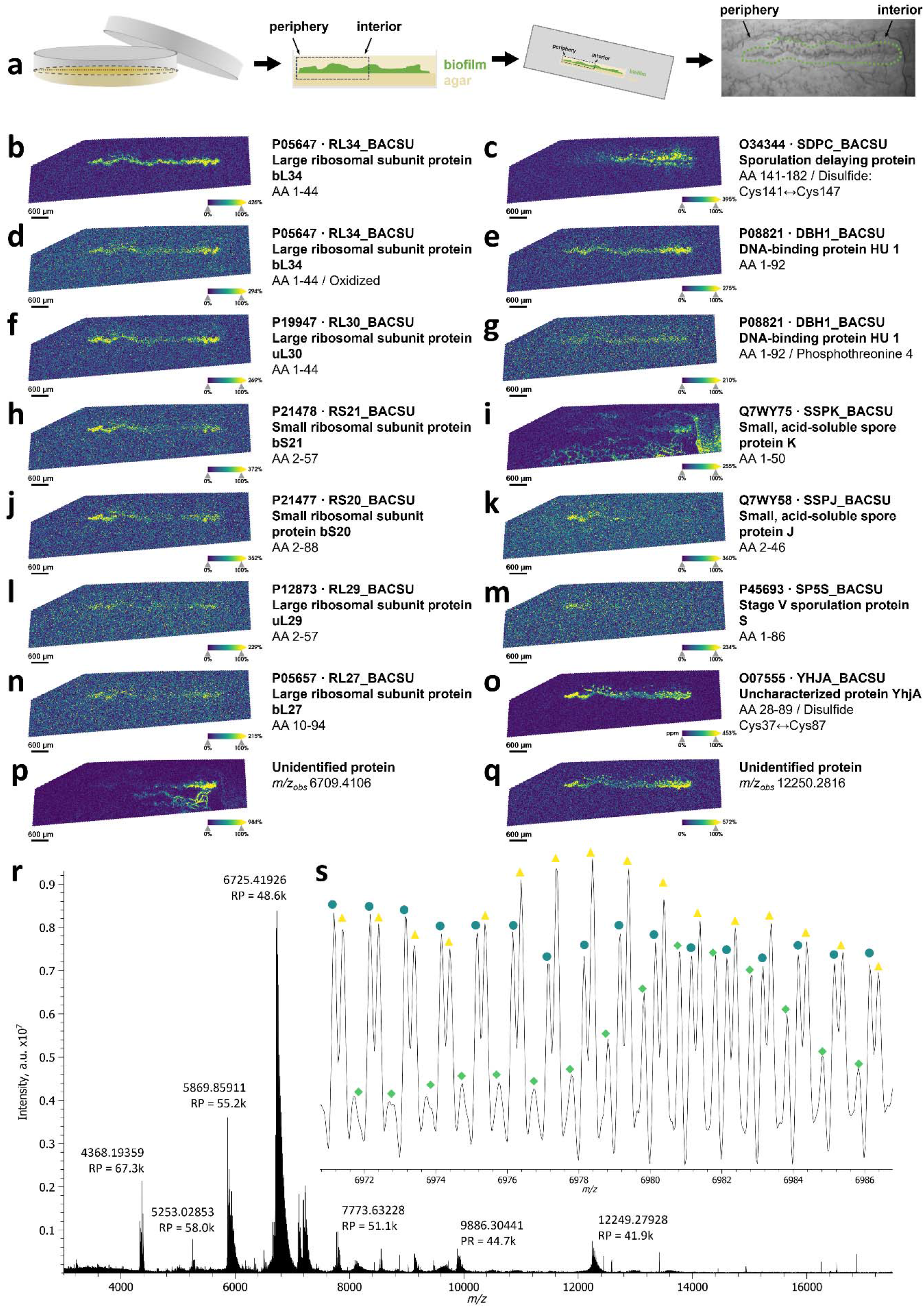
The sample preparation is outlined:**(a)** colonies grown on MSgg agar media were further embedded and cryosectioned at 20 µm as outlined in **Supplementary Information** and described in depth elsewhere.(2) After sample preparation, ion images are shown for each proteoform, the name, UniProt accession ID, and amino acid sequence with modifications (if any) are reported. Theoretical and experimental *m/z* values of the highest abundance isotope with calculated error can be found within **Supplementary Information. (b)** Large ribosomal subunit protein bL34 (P05647), **(c)** sporulation delaying protein (SDP; O34344), **(d)** oxidized large ribosomal subunit protein bL34 (P05647), **(e)** DNA-binding protein HU 1 (P08821), **(f)** large ribosomal subunit protein uL30 (P19947;), **(g)** phosphorylated DNA-binding protein HU 1 (P08821), **(h)** small ribosomal subunit protein bS21 (P21478;), **(i)** small acid-soluble spore protein K (Q7WY75), **(j)** small ribosomal subunit protein bS20 (P21477), **(k)** small acid-soluble spore protein J (Q7WY58), **(l)** large ribosomal subunit protein uL29 (P12873), **(m)** stage V sporulation protein S (P45693), **(n)** large ribosomal subunit protein bL27 (P05657), **(o)** a 7.1kDa protein, which is either uncharacterized protein YhjA (O07555) with a disulfide at Cys37↔Cys87 resulting in a [-2.01565 Da] shift, or regulatory protein DegR (P68731) with larger error, **(p)** a 6.1kDa unannotated protein, and **(q)** a 12.4kDa unannotated protein. The average spectrum from MALDI-MSI **(r)** is shown from *m/z* 3000 to 17500, with the complexity shown within an inset zoom **(s)** of the spectrum surrounding the base peak from *m/z* 6971 to 6987. Three distinct isotopic series from unidentified proteins are contained within this region, which are highlighted with sequential markers of teal circles, yellow triangles, and green diamonds. These series of isotopes are resulting from complex ion adduct substitutions and addition of PTMs. **SMART** annotation:(37) **S** (step size, spot size, total pixels) = 20 µm, 12 µm x 15 µm, 33084 pixels; **M** (molecular confidence) = MS1, <5 ppm; **A** (annotations) = MS1 matching from TDP LC-MS/MS; **R** (resolving power) = 48.6k at *m/z* 6725.4; **T** (time of acquisition) = 286 min.

Of the proteoform distributions, characteristic peptide toxins (e.g., SDP) are seen within the active interior of the biofilm **(Figure 1c)**,(26) while sporulating proteins **(Figure 1i, 1k, 1m)** and further uncharacterized proteins **(Figure 1o, 1q)** were found within the outer periphery. The presence of SDP **(Figure 1c)** in the central core of the colony could be indicative of cannibalistic cells recycling older generations, but the absence within the outer periphery and presence of characteristic sporulating proteins highly localized to the outermost periphery **(Figure 1i, 1k, 1m)**, suggests the biofilm is in early phases of sporulation. Stage V sporulation protein S could be overcome by other proteins or mutation,(27) but based on its colocalization with small acid-soluble spore (SASP) protein J **(Figure 1k)**, the presence of sporulated cells could be confirmed where SASPs are roughly 20% of the outer spore coat of *B. subtilis*. (28) These findings are supported by reports that cannibalistic toxins tend to prolong and delay sporulation,(13, 14) further demonstrating the utility of the detection of intact proteoforms for spatial subtyping of cell subpopulations.

TDP also detected sporulating delaying protein (SdpC) and immunity protein (SdpI) **(Supplementary Material)**, which offers immunity to cells through the sequestration of SDP.(29) These data demonstrate that tandem TDP of microbial colonies can complement MALDI-MSI analyses with coverage of larger proteoforms. When paired with laser capture microdissection (LCM), spatial/sub-sample specific proteoform information can be integrated readily.(30) Additionally, if paired with fluorescence microscopy, a robust multimodal pipeline to validate cell subpopulations and phenotypes is created. Several ribosomal proteins, including large subunits **(Figure 1b, 1f, 1l, 1n)** were also routinely annotated with initiator methionine truncations, and small subunits **(Figure 1h, 1j)** were also located throughout the main body of the biofilm. The broad detection of these ribosomal proteins lends credence to spatial biotyper analyses,(11) where speciation and localization of microbial cultures is feasible based on proteomic signatures. If cocultured colonies of bacteria and fungi are analyzed, or when working upon mutant knockouts or bioengineered strains, this information can be vital to understanding inter-microbial colonization and organization while further informing upon the cellular processes and phenotypes.

We identified many isotopic signatures that are resolved but separated by only a few hundred mDa such as the isotopic envelope series annotated by blue circles and yellow triangles **(Figure 1s)**. This complexity highlights the importance of sample-specific proteoform libraries and high mass-resolving power limiting this workflow to Fourier transform (FT)-MS instruments. While current applications were limited by incompatibility of ITO glass slides and LCM-TDP workflows,(30, 31) future works should leverage spatial LCM-TDP with MALDI-MSI for proteoform-informed spatial proteomics.(25) For example, several annotations would benefit from such serial analyses, such as the 7.1 kDa protein **(Figure 1q)** annotated as either uncharacterized protein YhjA (O07555) with a putative disulfide, or the regulatory protein DegR (P68731) with higher mass error. Protein YhjA possesses a signal peptide (AA 1-27) that is cleaved, and a high confidence disulfide link (C37↔C87) that was annotated due to the mass shift, and is supported by AlphaFold structural predictions **(Supplementary Figure 1)**, even though it is not listed within UniProt.(32, 33) Functionally, DegR stabilizes DegU phosphorylation with links to increased protease activity,(34) and regulates the transition of motile cells to sessile forms.(10) While the mass spectral error of this potential annotation is higher than current thresholds, this demonstrates the limits of high-resolution accurate mass isotopic pattern matching. In either case, this protein links major mechanisms producing “miner” cells that degrade extracellular biopolymers and peptides(35) and potential interactions with cannibalistic toxins that prolong sporulation.(14) It should be noted DegR was not detected by TDP, possibly due to analysis of a different culture or growth period, while other proteins such as DegV (36) were identified **(Supplementary Material)**. Overall, these results demonstrate that advanced measurement science is primed for future microbial applications and that further use of this instrumentation and pipeline can potentially unravel functions of many uncharacterized or understudied proteins with proper experimental design.

## CONCLUSIONS

This study demonstrates that proteoform-informed spatial proteomics can resolve the biochemical heterogeneity that governs microbial differentiation within *B. subtilis* biofilms. By integrating MALDI⍰MSI with sample-specific TDP libraries, we detected more than 285 proteoforms and revealed several distinct spatial distributions corresponding to cannibalistic, sporulating, and metabolically active subpopulations. These results show that proteoform⍰informed MSI can directly map functional microenvironments, offering mechanistic insight into the temporal progression of biofilm development. While these experiments were completed within a model system of *B. subtilis*, these methods for MALDI-MSI and TDP are universal and can be broadly applied to any microbial system that can be cultured on solid media. Additionally, while this is functionally limited to small proteoforms (< 20 kDa) for MALDI-MSI, the information gathered at fine spatial resolutions significantly enhances our understanding of the metabolic gradients and biomolecular mechanisms that shape community structure. The ability to localize peptide toxins, sporulation⍰associated proteins, ribosomal subunits, and uncharacterized proteins underscores the power of this approach to uncover both known and previously unappreciated contributors to microbial physiology.

## DECLARATIONS

### Conflicts ofInterest/Competing Interests

The authors declare no conflicts of interest or competing interests.

### Data availability statement

The data presented within this article is publicly available and was uploaded to MassIVE, partner repository with the MassIVE accession: MSV000100434.

## Acknowledgements

The authors would also like to acknowledge Dr. Mikhail Belov at Spectroglyph, LLC as well as Dr. Gordon Anderson and Chris Anderson at GAA Custom Electronics, LLC for technical support for the MALDI source. We would also like to acknowledge Drs. Tobias Wörner, Kyle Fort, Maria Reinhardt-Szyba, and Alexander Makarov of Thermo Fisher Scientific for technical guidance and licensing of the instrument. We thank Dr. Matthew E. Monroe of Pacific Northwest National Laboratory for his assistance in depositing the raw proteomic data onto MassIVE. Portions of Figures were aided by the use of BioRender.com.

## SUPPLEMENTARY INFORMATION

The following is provided as supplementary information for this article includes: (1) “Supplementary Information” outlining sample preparation, mass spectrometry methods, and data analysis, supplementary figure S1, and a supplementary table of highlighted annotations, and (2) “Supplementary Material” in an output table of the TDP results from both mass spectrometry instruments used.

