## Supplementary Information for "Characterization of the biofilm landscape of *Bacillus subtilis* by spatial microproteomics"

**CORRESPONDING AUTHOR(S):**

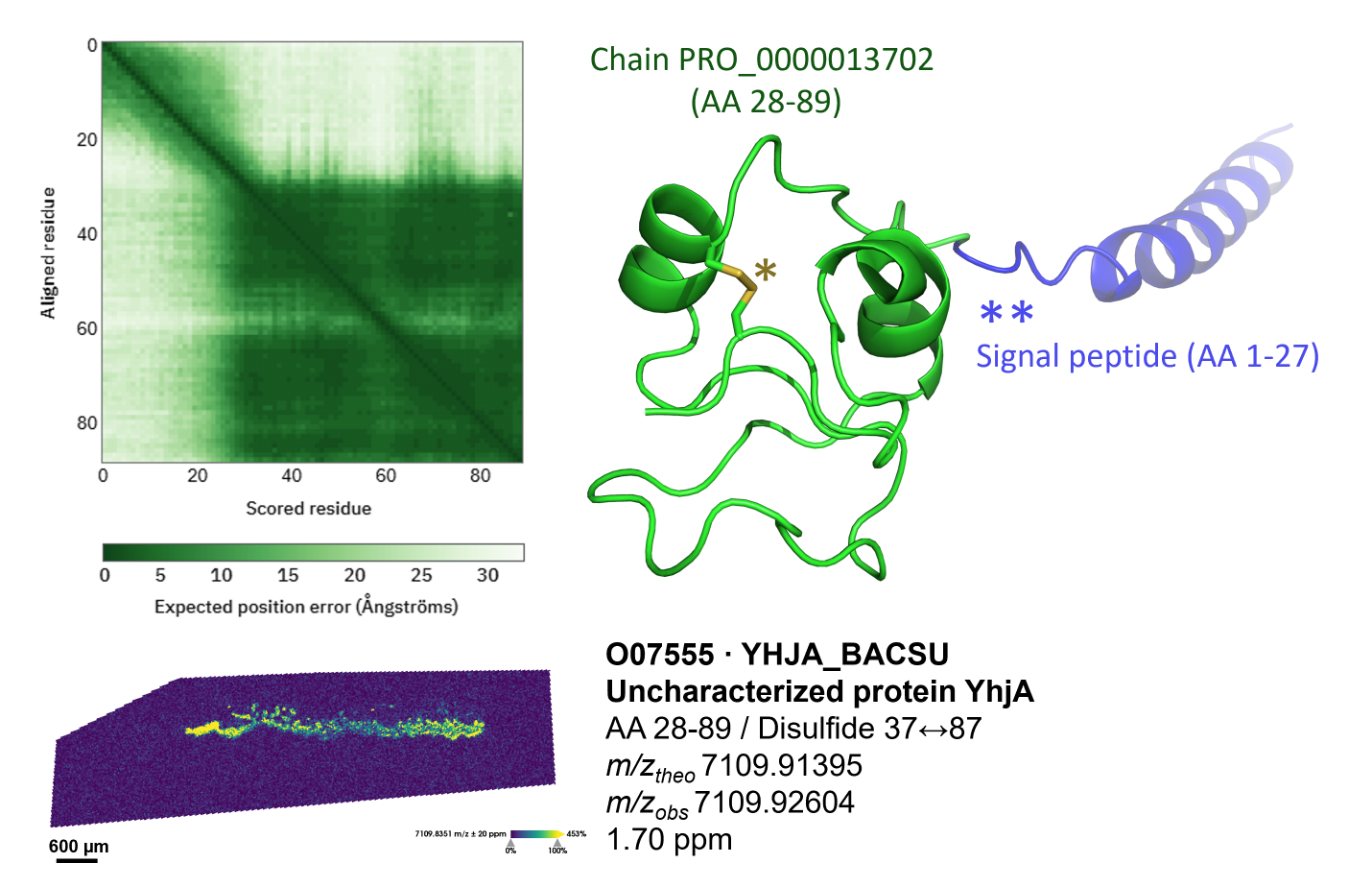

**Supplementary Figure S1:** Structural predictions of uncharacterized protein YhjA (O07555) are shown within AlphaFold2 dataset (AF-O07555-F1-v6) and were used to investigate the disulfide bond localization.(1, 2) The predicted aligned error (PAE) for confidence in the relative position of two residues is shown from AlphaFold2, where within the main chain (PRO_0000013702) corresponding to amino acids (AA) 28-89 and the signal peptide (AA 1-27)** are highlighted within green and blue. High confidence is expressed within the main chain, with a disulfide shown within yellow (*) between Cys37↔Cys87. While existence of the protein is inferred from homology within the reviewed UniProt database, based upon high resolution accurate mass matching, differential adduct formation (e.g., [M+H]^+^ and [M+Na-H]^+^), and AlphaFold2 predictions this annotation builds confidence and distributions throughout the biofilm are suggestive of being an protein integrated within the extracellular matrix (ECM) of *B. subtilis*.

| **Proteoform annotated** | **UniProt ID** | **Amino acid sequence** | **Post-translational modification (PTM,**  **if any present)** | ***m/z* theoretical** | ***m/z* observed** | **Calculated error (ppm)** |
| --- | --- | --- | --- | --- | --- | --- |
| Large ribosomal subunit protein bL34 | P05647 | 1-44 |  | 5253.02562 | 5253.02853 | 0.55 |
| Sporulation delaying protein | O34344 | 141-182 |  | 4336.20593 | 4336.20244 | -0.80 |
| Large ribosomal subunit protein bL34 | P05647 | 1-44 | Oxidized | 5269.02053 | 5269.01518 | -1.02 |
| DNA-binding protein HU 1 | P08821 | 1-92 |  | 9884.30074 | 9884.32167 | 2.11 |
| Large ribosomal subunit protein uL30 | P19947 | 2-59 |  | 6506.65523 | 6506.64602 | -1.42 |
| DNA-binding protein HU 1 | P08821 | 1-92 | *****Phosphorylated, Thr-4 | 9963.25924 | 9963.22323 | 3.16 |
| Small ribosomal subunit protein bS21 | P21478 | 2-57 |  | 6698.81197 | 6698.81030 | 0.24 |
| Small acid-soluble spore protein K | Q7WY75 | 1-50 |  | 5869.8826 | 5869.85911 | 4.01 |
| Small ribosomal subunit protein bS20 | P21477 | 2-88 |  | 9468.31450 | 9468.33166 | 1.81 |
| Small acid-soluble spore protein J | Q7WY58 | 2-46 |  | 5030.66695 | 5030.66146 | -1.09 |
| Large ribosomal subunit protein uL29 | P12873 | 1-66 |  | 7713.28033 | 7713.26573 | -1.89 |
| Stage V sporulation protein S | P45693 | 1-86 |  | 8796.82551 | 8796.83830 | 1.45 |
| Large ribosomal subunit protein bL27 | P05657 | 10-94 |  | 9207.91645 | 9207.91854 | 0.23 |
| ******Uncharacterized protein YhjA | O07555 | 28-89 | *******Unreduced disulfide, Cys37↔Cys87 | 7109.91395 | 7109.92604 | 1.70 |
| ******Regulatory protein DegR | P68731 | 1-60 |  | 7109.78424 | 7109.92604 | 7.15 |

**Supplementary Table S1:** Annotations of proteoform, UniProt IDs, PTM if present, the listed modification, the theoretical *m/z* of the proteoform, the observed *m/z* of the largest isotope of the proteoform, and the mass error is reported in ppm. *****Phosphotheonine is only reported on Thr-4 within UniProt for DNA-binding protein Hu 1, ******lists two potential proteoforms for one singular *m/z* signature, *******refers to a disulfide not reported within UniProt but is predicted by AlphaFold2 – see **Supplementary Figure S1**.
